# A single episode of CMV reactivation initiates long-term expansion and differentiation of the NK cell repertoire

**DOI:** 10.1101/2022.02.28.482225

**Authors:** Norfarazieda Hassan, Suzy Eldershaw, Christine Stephens, Francesca Kinsella, Charles Craddock, Ram Malladi, Jianmin Zuo, Paul Moss

## Abstract

NK cells play an important role in suppression of viral replication and are critical for effective control of persistent infections such as herpesviruses. Cytomegalovirus infection is associated with expansion of ‘adaptive-memory’ NK cells with a characteristic CD16^bright^CD56^dim^ NKG2C+ phenotype but the mechanisms by which this population is maintained remain uncertain. We studied NK cell reconstitution in patients undergoing haemopoietic stem cell transplantation and related this to CMV reactivation. NK cells expanded in the early post-transplant period but then remained stable in the absence of viral reactivation. However, a single episode of CMV reactivation led to a rapid and sustained 10-fold increase in NK cell number. NKG2C expression was increased on all NK subsets although the kinetics of expansion peaked at 6 months on immature CD56^bright^ cells whilst continuing to rise on the mature CD56^dim^ pool. Phenotypic maturation was observed by acquisition of CD57 and KIR expression. Transplantation from CMV-seropositive donors was associated with more rapid NK expansion and superior control of viral reactivation. Effective control of viral reactivation was seen when the peripheral NK cell count reached 20,000/ml. These data show that a single episode of CMV reactivation acts to reprogramme hemopoiesis to drive a sustained modulation and expansion of the NK cell pool and reveal further insight into long term regulation of the innate immune repertoire by infectious challenge.

**Author Summary:** im to highlight where your work fits within a broader context; present the significance or possible implications of your work simply and objectively; and avoid the use of acronyms and complex terminology wherever possible. The goal is to make your findings accessible to a wide audience that includes both scientists and non-scientists

Innate immune cells respond rapidly to infectious challenge and it has previously been thought that they were unable to ‘learn’ from specific infecitons but provided only short term control. However, in recent years it has become apparent that some subsets of innate cells, termed adaptive natural killer (NK) cells, are permanently increased in people who have become infected with cytomegalovirus. The mechanism by which this is maintained is not known. We studied the kinetics of adaptive NK cell expansion in patients who had recently undergone stem cell (bone marrow) transplantation as CMV very often ‘reactivates’ in this setting and allows study of how this affects the immune system. We find that one short term episode of viral replication led to a large and prolonged expansion of adaptive NK cells. Indeed, their levls increasd 11-fold over 10 months.

## Introduction

Natural killer (NK) cells have the capacity to mediate lysis of virally-infected and transformed cells through integration of signalling from activating and inhibitory receptors^1,2^ and play an important role in the control of herpesvirus infections^3^. Cytomegalovirus (CMV) is not cleared after initial infection but establishes a persistent infection which acts to shape the subsequent peripheral NK cell repertoire^4^. In particular, CMV-seropositive people develop expansion of a population of ‘adaptive NK cells’ with a predominant CD56^dim^CD16+NKG2C+CD57+ phenotype ^5-8^. This may be driven largely by engagement of NKG2C with viral peptides from the UL40 protein presented by HLA-E on virally infected cells^9^ although such populations are also observed in people with an NKG2C-/- genotype^10^. The NKG2 proteins are important regulators of NK activation and both NKG2A and NKG2C form a heterodimer with CD94 that binds to HLA-E^11,12^. However, whilst NKG2A acts as an inhibitory receptor and is predominantly a marker of immature NK cells, NKG2C is an activating receptor that is expressed on highly differentiated populations. The potential functional importance of this adaptive NK cell subset remains somewhat unclear although it has high capacity for antibody-directed cell cytotoxicity (ADCC).

Allogeneic haemopoietic stem cell transplantation (HSCT) is an important therapeutic modality within haemato-oncology and has the potential to cure patients with chemotherapy-resistant disease ^13^. Much of this curative effect is mediated through the activity of the donor immune system^14^ and there is increasing interest in the potential role of NK cells in this regard. The number of NK cells within the donor graft correlates with protection from disease relapse after allo-HSCT^15^ and early NK reconstitution can help to suppress GvHD whilst increasing the ‘graft versus leukaemia’ effect ^16,17,18^. Patients are highly immune suppressed in the early period after HSCT and reactivation of CMV is an important clinical complication that is associated with increased rates of morbidity and mortality ^19,20^. However, despite the increased short term clinical risk mediated by CMV reactivation it has also been demonstrated that viral reactivation may reduce subsequent post-transplant leukaemia relapse ^21,22,23^. The mechanism behind this observation remains uncertain but may relate to the generation of high numbers of CMV-driven donor NKG2C+ NK cells that increase capacity for control of host tumour cells. Although CMV reactivation after hempoietic or solid organ transplantation has been shown to boost the number of mature NKG2C+ NK cells ^24,25^ relatively little is known regarding the kinetics of this response from the time of reactivation. Here we determined the profile peripheral NK cell repertoire in patients after an episode of CMV reactivation following HSCT. Importantly, samples were taken immediately after viral reactivation and then followed prospectively. We observe that a single episode of viral reactivation leads to a long term expansion of NKG2C+ NK cells that transitions across the NK differentiation repertoire over a period of at least 10 months. Transient viral reactivation is thus seen to reprogramme long term NK cell differentiation and reveals further insights into pathogen-driven regulation of lymphopoiesis. me

## Results

### A single episode of CMV reactivation following HSCT drives rapid and long term expansion of NK cells

66 patients who underwent allo-HSCT with a 10/10 HLA-matched sibling or unrelated donor were recruited to this prospective study (Table 1) and PCR monitoring for CMV reactivation performed weekly. CMV reactivation was observed in 41 patients (62%) at an average time of 52 days post-transplant. Blood samples were then taken at the time of reactivation and at weeks 1, 2, 3, 4, and months 3, 6, 10, after CMV reactivation. Control post-transplant samples were also taken from patients without evidence of viral reactivation and were matched for equivalent timepoints of collection. Blood samples were also available pre-transplant and pre-reactivation from almost all donors.

**Table 1.** Demographics of patients with an episode of CMV reactivation following HSCT. MUD, Matched Unrelated Donor; AML, Acute Myeloid Leukaemia; ALL, Acute Lymphoblastic Leukaemia; MDS, Myelodysplastic syndrome; CMML, Chronic Myelomonocytic Leukaemia; NHL, Non-Hodgkin Lymphoma; RCMD, Refractory cytopenia with multilineage dysphasia; aGvHD, acute Graft versus Host Disease; cGvHD, chronic Graft versus Host Disease. Flu, fludarabine; Melph, melphalan; Campath, alemtuzumab; Bu, busulphan; ATG, anti-thymocyte globulin; Flamsa, fludarabine with amascrine and cytarabine; BEAM, BCNC, etoposide, cytarabine, melphalan; Cyclo, cyclophosphamide; TBI, total body irradiation; TLI, total lymphoid irradiation.

|  | Number<br>(n=41) | Percentage<br>(%) |
| --- | --- | --- |
| **Age**<br>20-70 years (median: 59 years) |  |  |
| **Average time of viral reactivation (days post transplant)**<br>52±5 (median: day 43)<br>Range (10—125 days) |  |  |
| **CMV serostatus (Recipient/Donor)** |  |  |
| R+/D+ | 22 | 54 |
| R+/D- | 15 | 37 |
| R-/D- | 3 | 7 |
| R+/D equivocal | 1 | 2 |
| **Sex** |  |  |
| Male | 24 | 58.5 |
| Female | 17 | 41.5 |
| **Donor type** |  |  |
| Sibling | 13 | 31.7 |
| Match unrelated | 25 | 61 |
| Related | 1 | 2.4 |
| Cord | 2 | 4.9 |
| **Diagnosis** |  |  |
| AML | 16 | 39 |
| ALL | 4 | 10 |
| MDS | 9 | 22 |
| Others (CMML, NHL, RCMD) | 12 | 29 |
| **Relapse incidence** |  |  |
| Relapse | 10 | 24 |
| Non-relapse | 31 | 76 |
| **GvHD** |  |  |
| aGvHD | 24 | 59 |
| cGvHD | 1 | 2 |
| aGvHD and cGvHD | 3 | 7 |
| no GvHD | 13 | 32 |
| **Conditioning regimen** |  |  |
| Flu/Melph/Campath | 15 | 35.6 |
| Flu/Bu/ATG | 7 | 17.1 |
| Flamsa/Bu | 8 | 19.5 |
| Beam/Campath | 5 | 12.2 |
| Others (Cyclo/Flu/TBI, Cyclo/TBI, TLI/ATG, Bu/Cyclo, Cyclo/TBI, Flu/beam/Campath) | 6 | 14.6 |
| **Number of reactivations** |  |  |
| Single viral episode | 22 | 54 |
| Multiple viral episodes | 19 | 46 |

In initial studies the percentage of NK cells within peripheral blood was determined using flow cytometry (Fig. 1). The proportion of NK cells was markedly higher in patients compared to healthy donors (HD) and reflects early and robust NK reconstitution in the early post-transplant setting. Indeed, NK cells comprised over 50% of PBMC within the first 4-weeks post-HSCT and prior to any episode of viral reactivation. This value remained stable following CMV reactivation before declining gradually after 4 weeks towards the normal range (Fig. 1A). Moreover, the absolute number of NK cells also increased rapidly following CMV reactivation but then continued to rise further over the next 10 months. Indeed, values rose 11-fold from 3 cells/μl at the time of reactivation to 35 cells/μl at the final timepoint of assessment at 10 months (Fig. 1C). In contrast, the NK cell number remained very stable in patients who did not experience CMV reactivation (Fig. 1D). These observations reveal that a single episode of CMV reactivation drives rapid and long term NK cell expansion.

**Figure 1.**
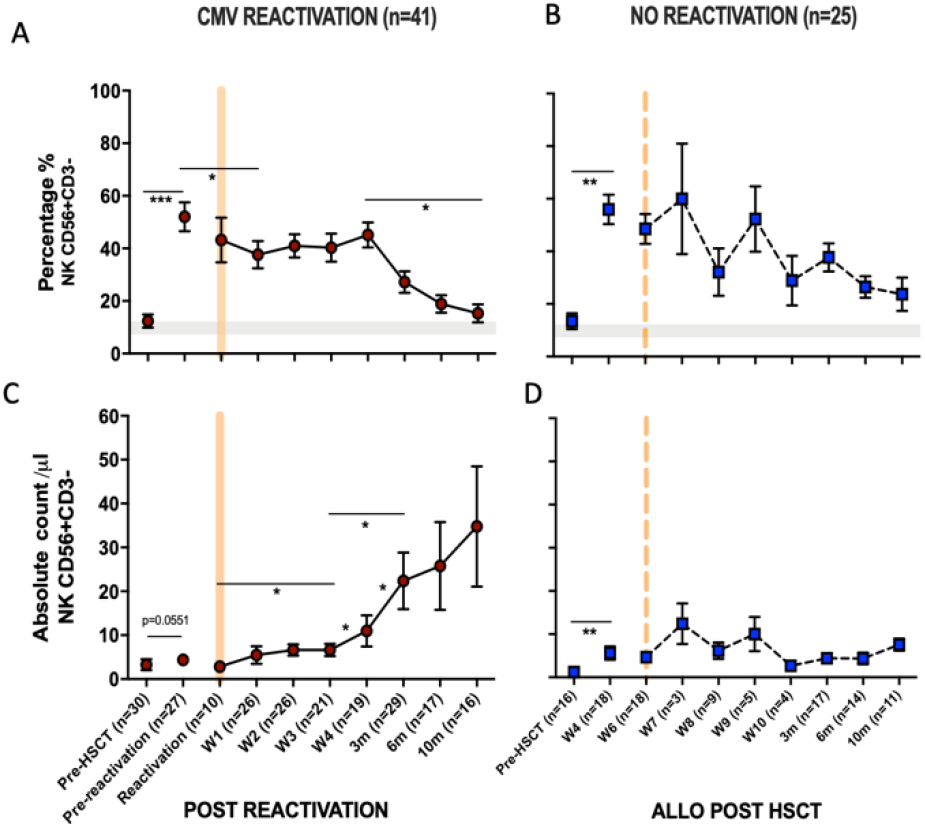
NK cells undergo long term expansion after a single episode of CMV reactivation. The percentage and number of peripheral NK cells was measured at regular timepoints prior to, and following, an episode of CMV reactivation in patients after HSCT. Matched timepoint samples were obtained from patients without an episode of reactivation. Sample times are shown as week (W) or month (m). (A) Percentage of NK cells in patients with CMV reactivation (B) Percentage of NK cells in patients without CMV reactivation (C) Number of NK cells in patients with CMV reactivation (D) Number of NK cells in patients without CMV reactivation Patients were classified as undergoing an episode of CMV reactivation when viral load reached > 200 copies/ml. Grey line represents the percentage range of NK cells within healthy donors.

### CD56^dim^ CD16^bright^ NK cells expressing NKG2C undergo a rapid and sustained expansion following CMV reactivation in patients after allo-HSCT

CD56^dim^ CD16^bright^ NK cells comprise the dominant peripheral population of NK cells and we next went on to assess the relative expression of NKG2C and NKG2A on this subset in relation to CMV reactivation (Fig. 2A). Average expression of NKG2C in healthy CMV seronegative donors was 1.4% (±0.2) compared to 11% (± 3.5%) in seropositive individuals (p<0.0001). Values for NKG2A^+^ were 50% (±2.6%) and 40% (±3.2%) respectively (p=0.0065) indicating that CMV-driven expansion of the NKG2C+ pool comes at the expense of a decrease in the NKG2A+ repertoire (Fig. 2B).

**Figure 2.**
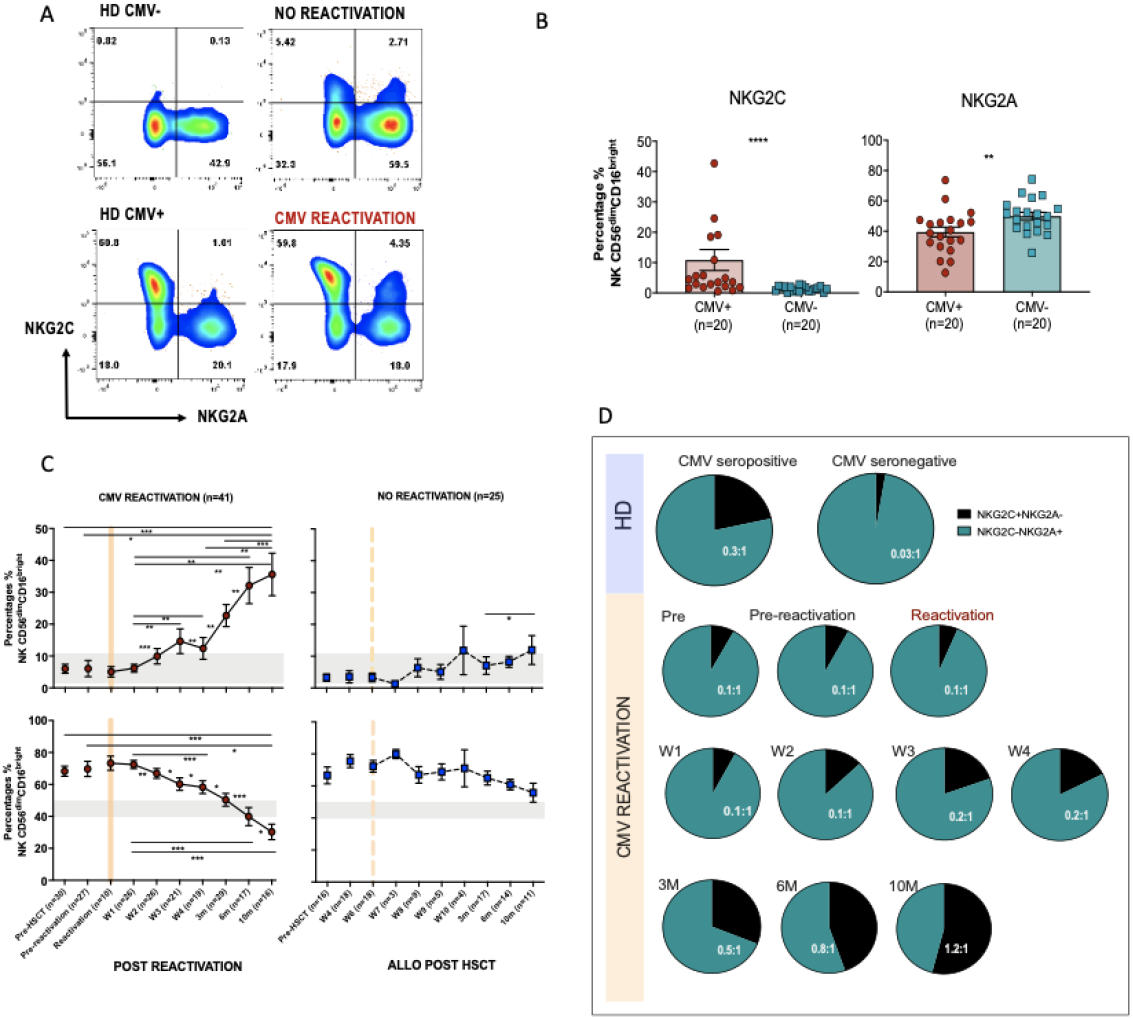
Rapid and long term expansion of the NKG2C+NKG2A-NK cell subset following CMV reactivation in patients after allo-HSCT. (A) Gating examples to show NKG2C/NKG2A expression on CD56^dim^CD16^bright^ NK subsets in CMV seropositive or seronegative healthy donors (HD) and allo-HSCT patients following CMV reactivation or no-reactivation. (B) Percentage of NKG2C+ and NKG2A+ NK cells within the CD56^dim^CD16^bright^ subset in CMV seronegative and seropositive HDs. (**p<0.01, ****p<0.001; Mann Whitney test). (C) Percentage of NKG2C+ (Top) and NKG2A+ (bottom) NK cells from CD56^dim^CD16^bright^ population in patients following reactivation (left) and without reactivation patients (right). Grey lines represent the range of NKG2C+ and NKG2A+ NK cells in HDs. (*p<0.05, **p<0.01, ***p<0.005; Wilcoxon sign rank test for matched time points). (D) Ratio of NKG2C+ and NKG2A+ cells within CD56^dim^CD16^bright^ population in HSCT patients with CMV reactivation and in healthy donors.

Within CMV-seropositive HSCT patients the percentage of NKG2C^+^ NK cells was comparable to that seen in CMV-seropositive controls prior to reactivation. However, a rapid expansion of CD56^dim^CD16^bright^ NK cells expressing NKG2C was observed within the first 7 days after viral reactivation. Furthermore, this population continued to expand for a further 10 months following clearance of viral reactivation such that the NKG2C^+^ cell proportion represented up to 36% of the total NK cell pool, a value far higher than the normal range in CMV-seropositive healthy donors (Fig. 2C). A much more modest increment was seen in patients who did not suffer an episode of CMV reactivation. The proportion of NKG2A^+^ cells within HSCT patients was initially higher than in healthy donors and is likely to reflect production of immature NKG2A^+^ NK cells during NK reconstitution. However, CMV reactivation accelerated NK cell maturation with an early decrease in the proportion of NKG2A^+^ cells after reactivation and a continuing fall to ~30% at month 10, a value below that seen in controls (Fig. 2C)

The NKG2C:NKG2A ratio was next determined on the NK cell pool at different time points. Within healthy donors this was very low at 0.03:1 in CMV-seronegative people compared to 0.3:1 in the CMV-seropositive group (Fig. 2D). Within HSCT patients the ratio was initially 0.1:1 but following viral reactivation it increased to reach 0.2-0.5:1 at 3 months, comparable to values seen in CMV-seropositive HD. Moreover, this increased further such that at 10 months the NKG2C+ cell percentage exceeded that of NKG2A+ with a ratio of 1.2:1. These data show that there is a rapid and long term expansion of the NKG2C+ NK cell subset following CMV reactivation in patients after allo-HSCT and that the relative proportion of this subset expands above that seen in healthy CMV seropositive people.

### NKG2C+ expression increases on all NK cell subsets following CMV reactivation and peaks earlier on immature populations

CMV-driven adaptive memory NK cells have been characterised previously only within the CD56^dim^CD16^bright^ NK cell subset. The relative expression of CD16 and CD56 is a marker of NK cell differentation and, given the finding of increased NKG2C expression on the cytotoxic CD56^dim^CD16^bright^ pool following CMV reactivation, we next went on to determine NKG2C and NKG2A expression on additional NK cell subsets (Fig. 3A).

**Figure 3.**
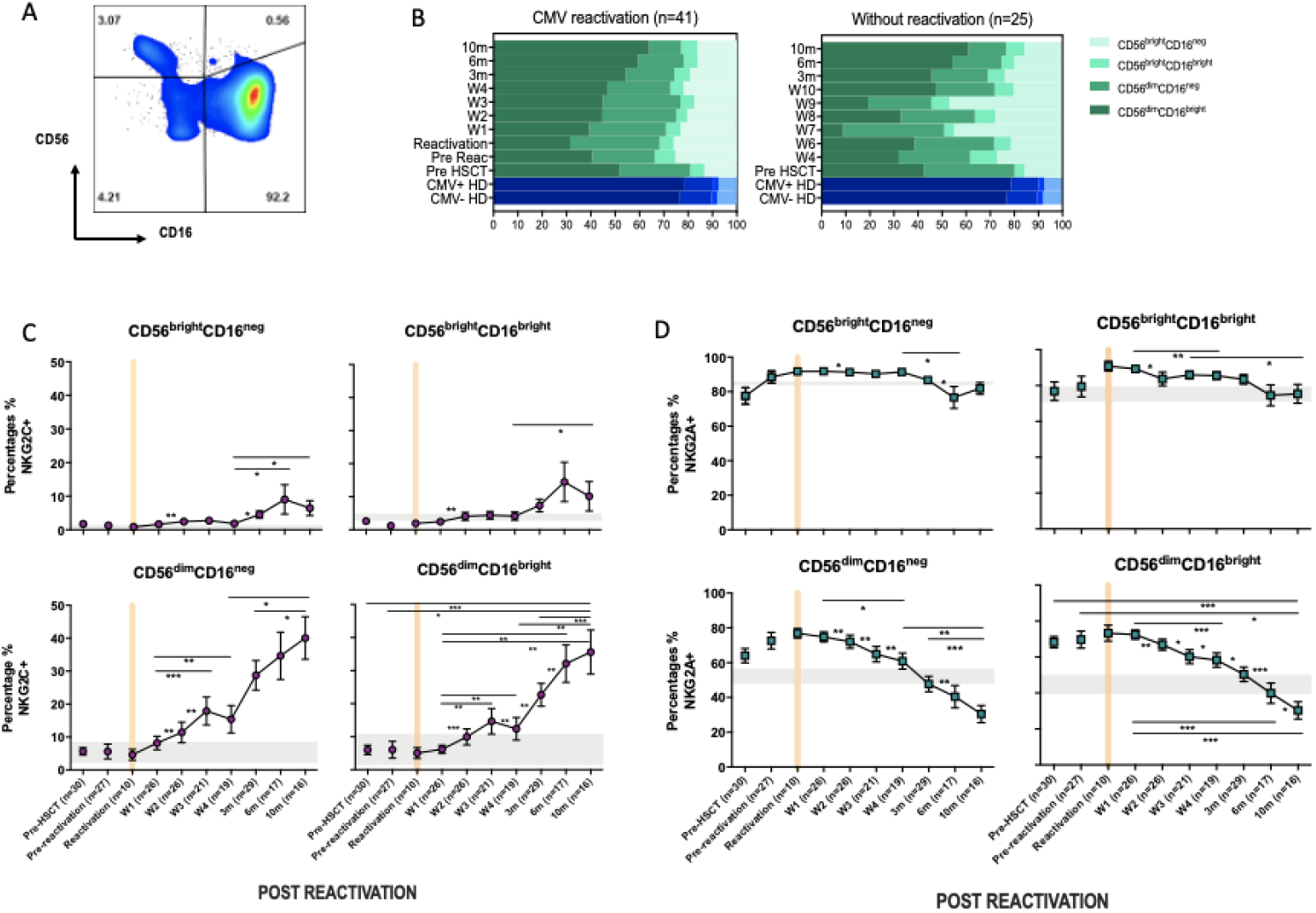
CMV reactivation leads to increased expression of NKG2C+ on all CD56^dim^ and CD56^bright^ subpopulations. (A) Gating to show separation of four NK subsets according to CD16 and CD56 expression. (B) Relative distribution of NK subsets in patients following CMV reactivation (left) and those without reactivation (right) (C) Relative NKG2C expression on NK subsets following CMV reactivation. (D) Relative NKG2A expression on NK subsets following CMV reactivation. Grey lines represent the range of NKG2C+ and NKG2A+ NK cells in HDs. (*p<0.05, **p<0.01, ***p<0.005; Wilcoxon sign rank test for matched time points).

The percentage of CD56^bright^ cells was increased in HSCT patients compared with controls, irrespective of CMV reactivation, and reflects accumulation of immature NK cells during early post-transplant lymphoid reconstitution (Fig. 3B). Interestingly, the expression of NKG2C^+^ increased on all NK subsets and was associated with reciprocal loss of NKG2A^+^ (Fig. 3C). NKG2C expression was 3-4 fold higher on mature cytotoxic CD56^dim^ NK subsets compared to CD56^bright^ cells. In addition, it was noteworthy that the kinetics of NKG2C expression differed between these two populations with NKG2C expression peaking at 6 months on CD56^bright^ cells whilst continuing to increase to month 10 on CD56^dim^ cells.

These data indicate that CMV-driven enhancement of NKG2C expression is imprinted at the immature stage of NK differentiation, peaking at 6-months, whilst continuing to accumulate on more mature subsets for a further 4 months.

### Transplantation from CMV seropositive donors is associated with more rapid expansion of NKG2C^+^ NK cells following CMV reactivation and superior control of viral replication

We next went on to determine how the CMV-serostatus of the haemopoietic transplant donor (D) acted to influence the magnitude of viral reactivation and profile of NK maturation following CMV reactivation in the patient (‘recipient’; R). Transplantation using a CMV-seropositive donor will infuse CMV-specific immune memory populations within the stem cell graft whilst immune cells from a CMV-seronegative donor will need to develop a primary CMV-specific immune response to control reactivation. 54% of patients who suffered an episode of CMV reactivation patients had R+/D+ CMV-serostatus whilst 37% had an R+/D-profile.

The magnitude and kinetics of viral load at reactivation was influenced markedly by donor CMV serostatus. Of note, the initial viral load at the time of CMV reactivation was higher in R+/D+ patients (2350 ±810 copies/ml vs 897 ±270 copies/ml in R+/D-patients; (p=0.05) (Fig. 4A). However, the subsequent peak viral load was considerably higher in the R+/D- cohort, indicating a role for transferred CMV-specific immunity in accelerating control of viral replication. 19 patients experienced two or more episodes of viraemia whilst 22 experienced only a single reactivation and this was seen exclusively within the R+/D- subgroup. Second peaks of viraemia occurred most commonly between weeks 2 and 4 and were typically higher than seen at initial reactivation. Viral load became virtually undetectable when the absolute NK count rose above 20/μl suggesting that this may represent a potential biomarker for effective control of viraemia.

**Figure 4.**
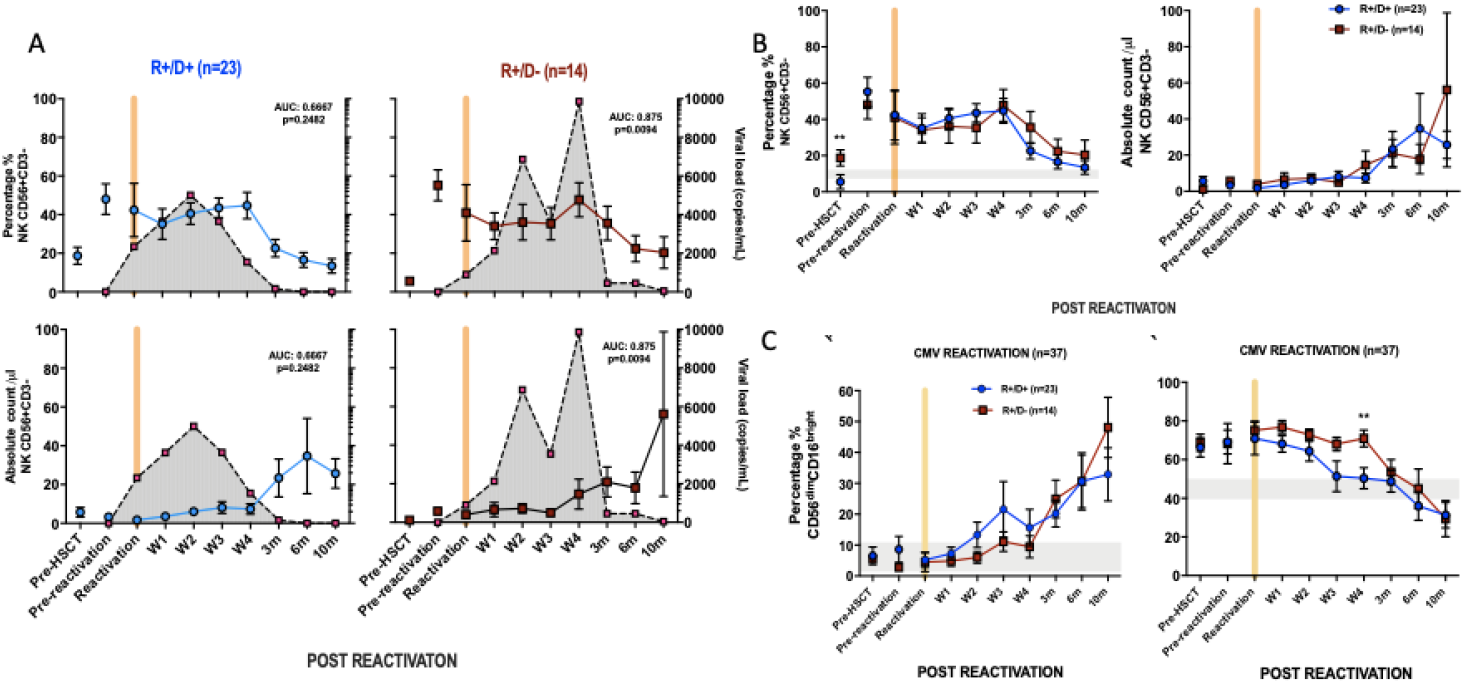
Influence of CMV serostatus of transplant donor on the kinetics of viral load at reactivation and subsequent profile of NK cells. NKG2C+ NKG2A-NK cells expansion showed delayed kinetic in R+/D- CMV risk groups. (A) Viral load and NK cell percentage and number following CMV reactivation in relation to CMV serostatus of donor (positive; D+ or negative; D-). Grey shading represents aggregate profile of CMV viral load over time. (B) Comparison of percentage and number of NK cells following CMV reactivation in relation to donor serostatus (R+/D+ in blue line and R+/D- in purple line). (C) Percentage of NKG2C+ (left) and NKG2A+ (right) NK cells following CMV reactivation in relation to donor serostatus (R+/D+ in blue line and R+/D- in purple line). Grey lines represent the range of NKG2C+ and NKG2A+ NK cells in healthy donors. (*p<0.05, **p<0.01, ***p<0.005; Wilcoxon sign rank test for matched phenotypes).

The kinetics of NK expansion varied according to donor serostatus and the percentage of NK cells was lower from W2 to W3 in R+/D- patients but then increased and rose to exceed the number in the R+/D+ group by month 10 (Fig. 4B). Reciprocal changes were observed in relation to expression of NKG2A (Fig. 4C)

These data suggest that transfer of memory NKG2C+ NK cells from a CMV seropositive donor contributes to accelerated control of viral reactivation whilst the delayed kinetics of NKG2C+ NK cell generation within R+/D- patients facilitates a larger peak viral load which ultimately drives development of a higher NK cell count in the longer term.

### NK cells show a more activated phenotype and produce higher cytokine levels after HCMV reactivation in HSCT patients

The pattern of expression of the early activation marker CD69 and the differentiation-associated protein CD57 was then assessed on the cytotoxic CD56^dim^CD16^bright^ NK subset. Of note, the percentage of CD69^+^ NK cells was much higher in HSCT patients compared to healthy donors at virtually all time points after transplant and is likely to reflect an influence of homeostatic proliferation (Fig. 5A). The highest level of expression was seen at the time of CMV reactivation and then fell over the next few weeks whilst values were stable in the absence of reactivation. The proportion of CD57^+^ NK cells decreased rapidly after CMV reactivation and remained low until week 4 before returning to the normal range at 3 months and increasing further thereafter (Fig. 5A). In contrast, the proportion of CD57+ NK cells remained very stable in the absence of reactivation.

**Figure 5.**
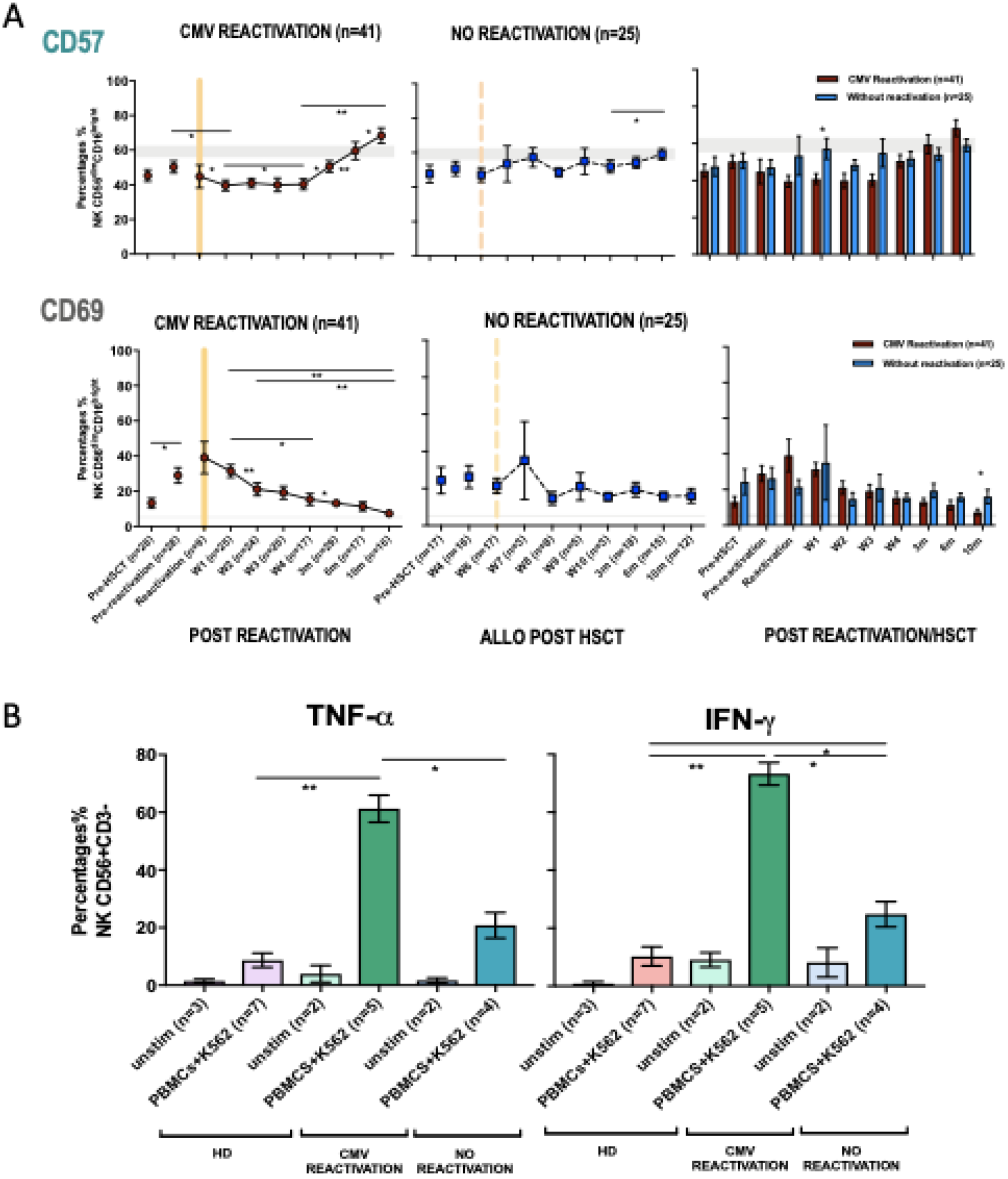
NK cells from HSCT patients who have undergone an episode of CMV reactivation demonstrate early proliferation with subsequent maturation and enhanced production of cytokines. (A) The percentage of CD57+ and CD69+ on NK cells from patients following CMV reactivation (purple line and bar) or without CMV reactivation (blue line and bar). Grey lines represent the range of NKG2C+ and NKG2A+ NK cells in healthy controls. (*p<0.05, **p<0.01, ***p<0.005; Wilcoxon sign rank test for matched phenotypes) (B) Enhanced IFN-γ and TNF-α production from NK cells of HSCT patients following prior CMV reactivation. Data are shown as percentage of cytokine-positive NK cells following K562 stimulation. (*p<0.05**p<0.01; Anova test).

The profile of TNF-α and IFN-γ production by NK cells was also determined in relation to prior history of viral reactivation. Due to limited sample availability, analysis was undertaken at 6-10 months post transplant and NK cells were co-cultured with K562 cells and assessed by intracellular flow cytometry. Higher baseline levels of cytokine production were seen posttransplant compared to controls and may reflect increased baseline proliferation as shown above (Fig. 5B). However, activation-induced cytokine production was markedly increased in patients who had experienced an episode of CMV reactivation. In particular, TNF-α and IFN-γ production was observed in 61% and 73% of NK cells respectively compared to only 21% and 25% in patients without reactivation (Fig. 5B). Cytotoxic responses against K562 were comparable between groups (data not shown).

These findings show that an episode of CMV reactivation serves to drive NK cell proliferation and differentiation and mediates a sustained increase in capacity for inflammatory cytokine production.

## Discussion

CMV establishes a persistent infection that is controlled by immune surveillance and leads to imprinting of the peripheral lymphoid repertoire^4^. We find that a single episode of viral reactivation leads to rapid and sustained expansion of NK subsets over a period of many months. Our studies were undertaken in immune suppressed patients after allogeneic stem cell transplantation but the findings are likely to inform directly on the mechanisms that control CMV reactivation within healthy immune competent people. As such they provide insight into the immune control of CMV replication and further demonstration of how pathogens can modulate the long term profile of innate immunity.

NK cells are amongst the first lymphocyte subsets to reconstitute after HSCT and this was reflected in both the high percentage of NK cells and an increased proportion of immature subsets in the early post-transplant period ^18^. However, an episode of CMV reactivation was seen to have a profound impact on the subsequent profile of NK cell maturation and expansion. CMV reactivation occurred at a median of 52 days which is consistent with previous reports and reflects early reactivation of virus from host tissues ^24^. The number of NK cells in the early post-transplant period was low but was not predictive of reactivation risk. In patients who did not experience an episode of viral reactivation the percentage and number of NK cells remained broadly stable over the next 10 months. In contrast, viral reactivation led to a significant expansion of NK cells within 3-weeks which then continued such that NK cell number increased by 11-fold after 10 months. Interestingly, this increase was accompanied by a reduction in the NK cell percentage within the peripheral repertoire and reflects the profound impact of CMV reactivation on expansion of other lymphoid subsets, most notably T cells ^26^. The nature of HSCT transplant conditioning and donor choice is likely to impact substantially on the profile of NK reconstitution and similar findings have been seen after umbilical donor transplantion ^24^ although an increase in NK number was not seen in the setting of haploidential transplantation ^27,28^.

A particular focus was on the kinetics of the adaptive memory NKG2+CD56^dim^CD16^high^ NK subset that is expanded in CMV seropositive donors. We took advantage of early sampling to show that this population increased within 2 weeks of reactivation and NKG2C expression increased incrementally from 4% to 35% of the CD56^dim^CD16^high^ pool over 10 months. Expression NKG2C or NKG2A was largely mutually exclusive and a comparable fall in the percentage of the NKG2A population was therefore seen over this period.

Previous studies have identified CMV-driven NKG2C+ adaptive memory cells only within the mature CD56^dim^ NK population but here we also found this to develop on CD56^bright^ populations. Indeed, a relative increase in NKG2C expression was seen on all four NK subsets as defined by CD56 and CD16 expression although the relative kinetics and magnitude were markedly different. The increment in NKG2C expression on the immature CD56^bright^ subsets was much more modest than seen on CD56^dim^ cells, with maximal values of 8% and 15% for the CD16^neg^ and CD16^bright^ subsets respectively. A particular finding of interest was that NKG2C^+^ expression on CD56^bright^ cells peaked at 6 months whilst that on the CD56^dim^ subset continued to increase at month 10. CD56^bright^ cells are believed to be precursors of the CD56^dim^ population ^29^ and as such these data indicate that NKG2C^+^ CD56^dim^ cell expansion following CMV reactivation is at least partially supported by differentiation from NKG2C+ CD56^bright^ NK cell precursors during the first 6-months (Fig. 6). However, studies in patients with *GATA2* mutation or PNH have indicated that adaptive cells may have self-renewing properties and as such the lineage derivation of this pool remains unclear ^30,31^. In addition recent data from murine models has shown that an ILC1-like splenic-resident memory pool is induced following murine CMV infection ^32^ and the broad impact of viral reactivation on the repertoire of innate memory is therefore likely to have been underestimated ^33^.

**Figure 6.**
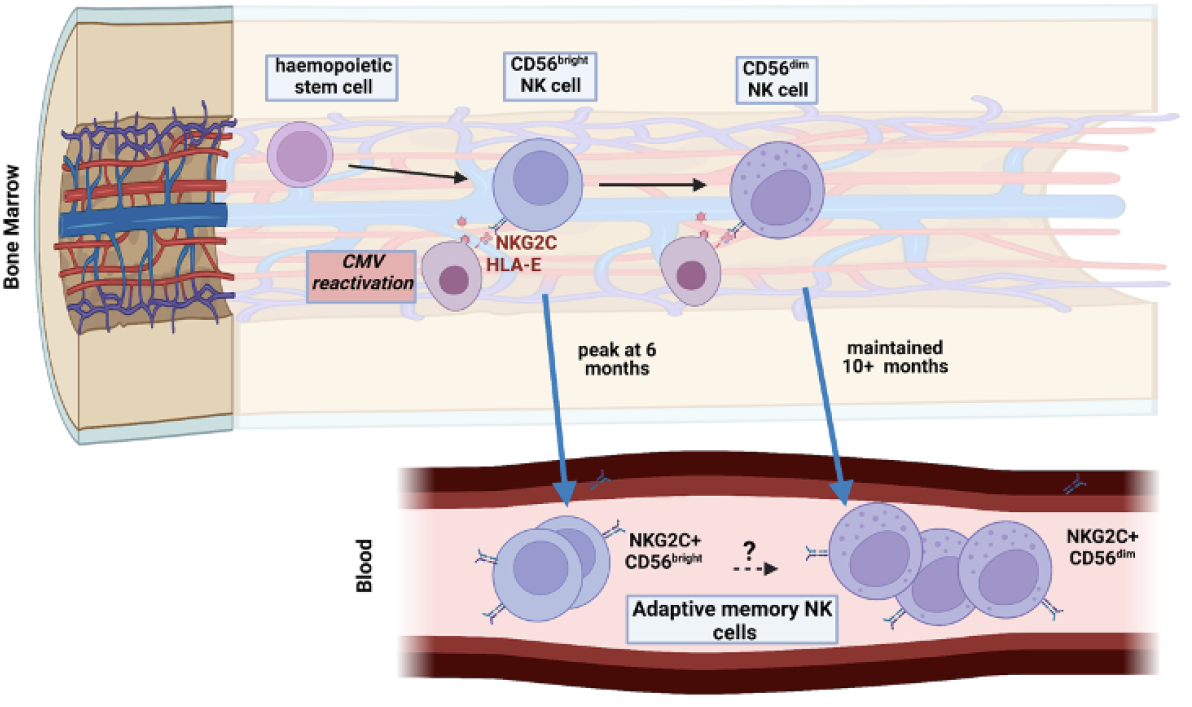
Model of maintenance of the adaptive memory NK cell pool through intermittent reactivation of CMV within bone marrow

The functional importance of transfer of ‘immune’ adaptive memory NK cells from the transplant donor was assessed by comparison of immune reconstitution between transplants with either a CMV-seropositive or seronegative donor. It was noteworthy that the initial viral load was elevated following transplantation from a CMV seropositive donor and this may reflect either viral replication from donor leucocytes or a consequence of donor CMV-specific immunity acting to transiently enhance short term viral load at reactivation ^34^. Despite this effect, peak viral load and longer term control of viremia were markedly improved following transplantation from a CMV-seropositive donor and recurrent episodes of reactivation were seen only with the use of a CMV-seronegative donor. NKG2C^+^ cell expansion was more rapid within the first four weeks after reactivation within the R+/D+ group, indicating that NK cells retain and transfer immune memory to facilitate control of viral reactivation ^35^. Furthermore, prospective analysis allowed assessment of NK number in relation to control of viral reactivation and identified that viral load was controlled in all subjects when the peripheral NK cell count reached 20,000/ml (20/μl), comparable to values of 18/μl associated with higher overall survival at day 60 post transplant^36^. The lymphocyte count in peripheral blood of healthy donors is ~10^6^/ml and, with an NK cell percentage of 2-10%, this number is comparable to the value of 20-100,000 NK cells/ml in CMV-seropositive healthy donors. Sustained control of viral reactivation is likely to also require T cells and the role of adaptive memory cells in antigen presentation is of note in this regard^37^.

Analysis of the kinetics of the extended phenotype of CD56^dim^CD16^high^ NK cells after viral reactivation focussed on CD57 and CD69. Expression of CD57, a marker of late differentiation, was somewhat lower in the first month post-reactivation but then increased continuously over subsequent months. As such, the classical expression of CD57 and KIR on adaptive memory NK cells is seen to develop late after reactivation^38^ and supports evidence for coupling of differentiation and proliferation of NK cells over an extended period^39,40,28^. It is not clear why expression was reduced during the first month after reactivation although this may reflect preferential recruitment of this subset into tissue. CD69 is upregulated upon NK cell activation^41^ and expression was broadly increased within all patients which is likely to reflect in the impact of homeostatic proliferation and immune reconstitution. However, values were at their highest level at the time of viral reactivation and then gradually fell.

The relative functional capacity of NK cells was assessed by the profile of cytokine secretion and cytotoxic capacity following activation *in vitro*. Cytokine production was generally enhanced within NK cells from HSCT patients compared to healthy donors and may reflect the increased proliferative status in this setting. However, CMV reactivation further primed for the production of high levels of TNF-α and IFN-γ, as seen previously^24^, and consistent with epigenetic modification of the *IFNG* locus ^42^. There is interest in the potential beneficial impact that may arise from CMV-driven enhancement of NK maturation ^43^ and reactivation has been associated with a reduction in the risk of leukaemia relapse in some studies although the underlying mechanisms remain unclear ^44,45^..

Homeostatic expansion^46,47^ and the high level of viral reactivation are both likely to enhance NK cell expansion in the post-transplant period and likely to underlie the ‘supra-physiological’ expansion of the NKG2C+ NK cell subsets that was observed. However, the broad features relating to the kinetics and phenotypic profile of viral-induced NK expansion are likely to be applicable to immune competent populations ^48^. NKG2C+ cells remain elevated for decades following CMV infection and, as hemopoietic and myeloid precursors are reservoirs for CMV latency, these observations suggest that this may be maintained by intermittent viral reactivation acting to modulate NK cell differentiation within the bone marrow. The finding that NK cell expansion is sustained for at least 10 months might appear surprising given that innate responses are typically regarded as reactive and short term. Very low levels of subclinical viral reactivation in the post-transplant setting cannot be ruled out but were not detected by sensitive PCR. Moreover, the findings concur with observations that other innate cells, such as neutrophils and monocytes, also show long term peripheral changes after acute events and suggest that epigenetic regulation of stem cell precursors acts to establish long term adaptive memory^49-50^. A limitation of our study was the cessation of sampling at 10 months after reactivation.

In conclusion we show that an episode of CMV reactivation drives rapid and long term expansion of NKG2C+ NK cells over a period of at least 10 months. These findings are of potential importance for management of CMV reactivation after HSCT and suggest that NKG2C+ NK cell monitoring might be used to predict immune capacity to control reactivation. Furthermore, this profile of expansion following a single episode of viral reactivation extends insight into pathogen-driven haemopoietic programming of stem cell precursors to enhance innate cell memory.

## Materials and Methods

### Patients and control subjects

41 patients who had undergone allo-HSCT in the Queen Elizabeth Hospital were recruited at the time of CMV reactivation following written informed consent under the BOOST ethics (RG 13-172) according to Declaration of Helsinki. 60ml of peripheral blood in sodium heparin (BD Vacutainer®, Cat#367876) was collected for the first four weeks following CMV reactivation and 36ml of heparinised and clotted blood was also available pre-transplant, pre-reactivation and for up to 10 months post-transplant. Samples were also collected from patients undergoing allo-HSCT but without CMV reactivation (n=26) at equivalent time points pre- and post-transplant for use as controls. This group of patients were also followed for up to 10 months post-transplant.

### Assessment of CMV serostatus and CMV reactivation

Patients were routinely monitored for CMV reactivation by quantitative PCR and reactivation was diagnosed by value of >200 copies/ml of CMV DNA in whole blood. CMV viral load was monitored for each week following reactivation. Patients were classified according to their CMV serostatus pair; recipient positive and donor positive (R+/D+); recipient positive and donor negative (R+/D-); recipient negative and donor positive (R-/D+) and recipient negative and donor negative (R-/D-). Donors with ambiguous CMV serostatus are classified as equivocal (R+/Dequi).

All patients received anti-viral propylaxis with acyclovir. Patients with a matched unrelated donor received IV aciclovir 500mg/m^2^ tds until mucositis resolved and then 800mg qds orally until day +100. Those who received a stem cell graft from a sibling donor were given oral aciclovir 200mg qds until day +35. Following viral reactivation patients received valganciclovir when viral load was >10,000 copies/ml.

### Healthy donors

40 healthy donors (HD) were used as controls following written informed consent (REC reference Number: 14/WM/1254). Up to 60 ml of peripheral blood was collected in sodium heparin tubes and CMV serostatus was determined.

### Isolation of Peripheral Blood Mononuclear Cells (PBMCs)

PBMCs were isolated by density gradient centrifugation over Lymphoprep™ (Stem cell Technologies, Cat #07861) within 24 hours of collection. Peripheral blood was diluted 1:1 with RPMI-1640 Medium (Sigma, Cat#R8758) with 1% Penicillin/Streptomycin (Gibco™ Cat#15070063), layered at a ratio of 2:1 and centrifuged at 2000 x g for 25 minutes (brake off). Buffy coats were harvested and resuspended in RPMI media and centrifuged at 500 x g for 10 minutes and the number of cells determined using a haemocytometer (Fast-Read 102®, Kova International, Cat #88010). Cells were washed and centrifuged at 500 x g for 10 minutes before being cryopreserved in freezing media (RPMI-1640 Medium [Sigma, Cat#R8758], 10% DMSO [Sigma, Cat #D2650]).

### Flow Cytometry analysis

Frozen cells were recovered by rapid thawing at 37°C in a water bath and centrifuging in prewarmed growth media at 500 x g for 10 minutes. PBMCs were surface stained with series of antibodies (anti-human CD158 [KIR2DL1/S1/S3/S5]-FITC [BioLegend, Cat#339504], anti-human CD158b [KIR2DL2/L3, NKAT2]-FITC [BioLegend, Cat#312604], anti-human CD158e1 [KIR3DL1, NKB1]-FITC [BioLegend, Cat#312706], anti-human NKG2C/CD159c-PE [R&D Biosystems, Cat#FAB138P], anti-CD14-ECD [Beckman Coulter, Cat#IM2707U], anti-CD19-ECD [Beckman Coulter, Cat#A07770], anti-CD3-PerCP-Cy5.5 [Beckman Coulter, Cat#300328], anti-human CD279 (PD-1)-PE-Cy7 (BioLegend, Cat#329916), anti-CD159a (NKG2A)-APC [Miltenyi Biotec; Cat#130-098-8123], anti-human CD56 (NCAM)-APC/Cy7 [BioLegend, Cat#318332], anti-KLRG1-APC-Vio770 [Miltenyi Biotec, Cat#322316], anti-human CD57-PB (BioLegend, Cat#322316], anti-human CD16-V500 [BD Biosciences, Cat#561314], anti-human CD69-PE-Cy7 [BioLegend, Cat#310190]). 5×10^5^ to 1×10^6^ cells were re-suspended in residual buffer and antibody panels added and incubated on ice for 20 to 30 minutes, then being washed in MACS buffer (1xPBS [Oxoid™/Thermo Scientific™ Cat#BR0014G], 0.5% BSA [Sigma, Cat#A7906] and 2mM EDTA [Sigma Cat#S8045] and centrifuged at 500 x g for 5 minutes. 1.5 μl of PI was also added and samples were run on the Gallios™ Flow Cytometer (Beckman Coulter, Inc). Data was analysed offline using Kaluza^®^ Flow Analysis software version 2.1a (Beckman Coulter Inc.) or FlowJo® v10 (TreeStar Inc). Flow cytometric gating for identification of NK cells was done according to methods and protocols established within the research group (Supplementary Figure 5).

### Statistical analysis

Data were analysed by using Mann-Whitney test for comparison between patients’ groups and Wilcoxon sign rank test for different time-points within similar patients. Friedman test was used for comparison in more than two groups within similar group of patients. All sample sizes in the different time-points are small and treated as not normally distributed and analysed by using GraphPad Prism 8 and SPSS 20.

## Acknowledgment

This study was funded by Bloodwise (Grant Code: 12052), Medical Research Council and Ministry of Education Malaysia.

## Author Contributions

PM, NH, JZ wrote the manuscript. PM, JZ designed the study. FK, CC, RM recruited participants. NH, SE, CS, JZ performed the experiments. NH, JZ analysed the data. All authors commented on the manuscript.

